# Developmental stage-specific changes in protein synthesis differentially sensitize hematopoietic stem cells and erythroid progenitors to impaired ribosome biogenesis

**DOI:** 10.1101/2020.08.22.262816

**Authors:** Jeffrey A. Magee, Robert A.J. Signer

## Abstract

Ribosomopathies encompass a collection of human genetic disorders that often arise from mutations in ribosomal proteins or ribosome biogenesis factors. Despite ubiquitous requirement of ribosomes for protein synthesis, ribosomopathies present with tissue- and cell-type-specific disorders, and blood is particularly affected. Several ribosomopathies present with congenital anemias and bone marrow failure, and accordingly, erythroid lineage cells and hematopoietic stem cells (HSCs) are preferentially impaired by ribosomal dysfunction. However, the factors that influence this cell-type-specific sensitivity are incompletely understood. Here, we show that protein synthesis rates change during HSC and erythroid progenitor ontogeny. Fetal HSCs exhibit significantly higher protein synthesis than adult HSCs. Despite protein synthesis differences, reconstituting activity of both fetal and adult HSCs is severely disrupted by a ribosomal mutation (*Rpl24*^Bst/+^). In contrast, fetal erythroid lineage progenitors exhibit significantly lower protein synthesis than their adult counterparts. Protein synthesis declines during erythroid differentiation, but the decline starts earlier in fetal differentiation than in adults. Strikingly, the *Rpl24*^Bst/+^ mutation impairs fetal, but not adult erythropoiesis, by impairing proliferation at fetal erythroid progenitor stages with the lowest protein synthesis relative to their adult counterparts. Thus, developmental and cell-type-specific changes in protein synthesis can sensitize hematopoietic cells to impaired ribosome biogenesis.

**Key Points:**

- Fetal HSCs synthesize much more protein per hour than young adult HSCs in vivo
- Fetal erythroid progenitors synthesize much less protein per hour than young adult erythroid progenitors in vivo
- Differences in protein synthesis dynamics distinguish fetal and adult erythroid differentiation
- A ribosomal mutation that reduces protein synthesis impairs fetal and adult HSCs
- Reduced protein synthesis impairs fetal but not adult erythroid progenitors

## Introduction

Adult hematopoietic stem cells (HSCs) maintain exquisite control of protein synthesis^1^. Inappropriately high or low protein synthesis rates impair HSC self-renewal and can lead to leukemia or bone marrow failure^1,2^. Adult HSCs exhibit very low protein synthesis compared to other hematopoietic cells^1,3^, which is required to protect HSCs from stress associated with protein misfolding^4^. Most adult HSCs are quiescent, but when forced into cycle, their protein synthesis remains low^1,3^. This raises the question of whether HSCs maintain low protein synthesis when they cycle in other contexts, such as fetal development^5^. Ontogeny-driven changes in protein synthesis carry potential implications for human disease. Ribosomopathies, such as Diamond Blackfan Anemia and Shwachman-Diamond Syndrome, cause congenital anemia and bone marrow (BM) failure, and present early in life^6-9^. This could reflect the germline nature of the mutations, but it could also reflect exquisite sensitivity to altered protein synthesis in fetal HSCs or erythroid progenitors. We therefore investigated whether hematopoietic stem and progenitor cells (HSPCs) undergo cell-type- and developmental stage-specific changes in protein synthesis that can alter the susceptibility to the deleterious effects of a ribosomal mutation *in vivo*.

## Methods

### Mice

*Rpl24*^Bst/+^ mice are previously described^1,10^. Transplant recipients were irradiated (540 rad x2) and transplanted with 10 fetal CD150^+^CD48^-^Lineage^-^Sca1^+^cKit^+^ HSCs^11^ and 3×10^5^ recipient- type BM cells, or with 5×10^5^ donor and 5×10^5^ recipient-type BM cells. Animals were housed in the Animal Resource Center at UT Southwestern or the UC San Diego Moores Cancer Center vivarium. Protocols were approved by the UT Southwestern and UC San Diego Institutional Animal Care and Use Committees.

### Cell isolation and flow cytometry

Cells were isolated by flushing long bones or crushing fetal livers in HBSS with 2% heat-inactivated bovine serum. Flow cytometric analysis/isolation were performed as previously described^4^.

### Protein synthesis

O-propargyl-puromycin^12^ (OP-Puro; Medchem Source) was injected into young adult or pregnant mice and analyzed as previously described^13^.

### Proliferation

2mg of EdU (Thermo) was injected into pregnant mice. Fetal livers were harvested 1h later, and EdU incorporation was assessed as previously described^3^.

## Results and Discussion

To investigate protein synthesis in adult and fetal HSPCs, we administered OP-Puro^1,12,13^ to young adult (2-3 month old) or timed pregnant mice. One hour later, we quantified OP-Puro incorporation in BM and E15.5 fetal liver HSPCs by flow cytometry. Consistent with previous reports^1,3^, adult HSCs^14^ exhibited significantly lower protein synthesis than restricted progenitors (Fig. 1A). In contrast, fetal HSCs did not exhibit low protein synthesis compared to most fetal progenitors (Fig. 1B). Fetal HSCs synthesized ∼2.6-fold less protein than CD127^-^Lineage^-^ cKit^+^Sca1^-^ myeloid progenitors^15^, but ∼1.3-1.6-fold more protein than unfractionated liver cells, erythroid progenitors, B cell progenitors and granulocytes (Fig. 1B). Because of differences in OP-Puro perfusion/uptake in distinct tissues^13^, we could not directly compare OP-Puro fluorescence between adult BM and fetal liver cells. Thus, we compared adult and fetal HSC protein synthesis by normalizing OP-Puro fluorescence to unfractionated BM and fetal liver cells or to Gr1^+^ cells. On these bases, fetal HSCs exhibited ∼3.5-4.5-fold higher protein synthesis than adult HSCs *in vivo* (Fig. 1C,D). Protein synthesis differences between adult and fetal HSCs cannot be fully explained by differences in proliferation, as adult HSCs driven to undergo rapid proliferation exhibit only modestly elevated protein synthesis^1,3^. Thus, protein synthesis rates change during HSC ontogeny, and they decline between fetal development and adulthood.

**Figure 1.**
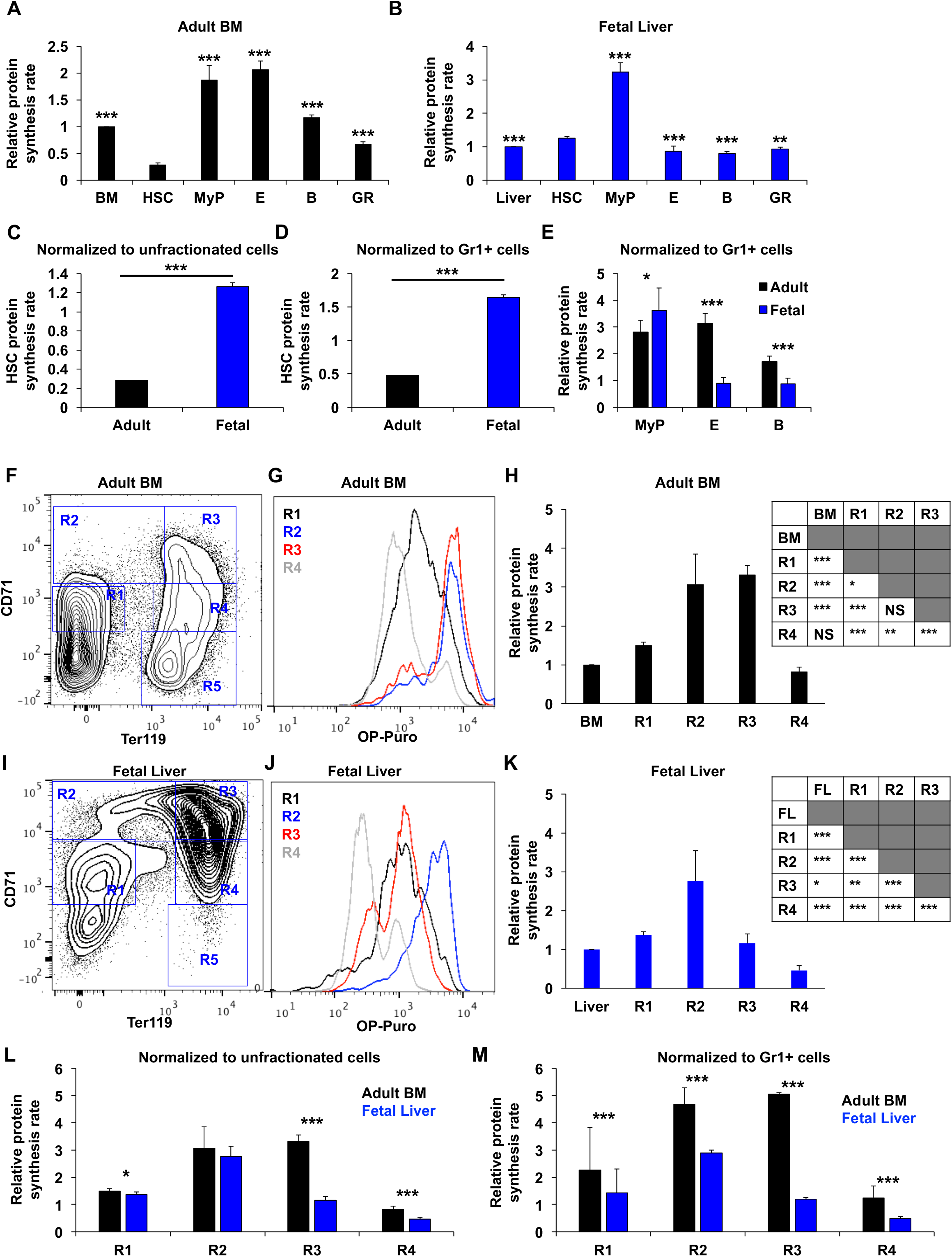
Distinct protein synthesis rates in fetal and adult hematopoietic stem and progenitor cells. (A,B) Relative protein synthesis in hematopoietic stem and progenitor cells relative to unfractionated cells in (A) young adult BM and (B) E15.5 fetal liver. Data are shown for unfractionated BM cells, liver cells, CD150^+^CD48^-^Lineage^-^Sca1^+^cKit^+^ HSCs, CD127^-^Lineage^-^ Sca1^-^cKit^+^ myeloid progenitors (MyP), CD71^+^Ter119^+^ erythroid progenitors (E), IgM^-^B220^+^ B lineage progenitors (B) and Gr1^+^ cells (GR). (C,D) Relative protein synthesis in young adult BM and fetal liver HSCs normalized to (C) unfractionated cells or (D) Gr1^+^ cells. (E) Relative protein synthesis in young adult BM and fetal liver restricted hematopoietic progenitor cells normalized to Gr1^+^ cells. (F) Representative flow cytometry plot showing gating strategy for R1-R5 erythroid lineage cells in young adult BM. (G) Representative histograms showing OP-Puro incorporation in R1-R4 erythroid progenitors in young adult BM. (H) Relative protein synthesis based on OP-Puro incorporation in young adult R1-R4 erythroid progenitors relative to unfractionated BM cells in vivo. Statistical differences are summarized in the adjacent table. (I) Representative flow cytometry plot showing gating strategy for R1-R5 erythroid lineage cells in E15.5 fetal liver. (J) Representative histograms showing OP-Puro incorporation in R1-R4 erythroid progenitors in E15.5 fetal liver. (K) Relative protein synthesis based on OP-Puro incorporation in E15.5 fetal liver (FL) R1-R4 erythroid progenitors relative to unfractionated liver cells in vivo. Statistical differences are summarized in the adjacent table. (L,M) Relative protein synthesis in young adult BM and E15.5 fetal liver R1-R4 erythroid progenitors normalized to (L) unfractionated cells or (M) Gr1^+^ cells. All data represent mean ± standard deviation (SD). N=7 adult mice and 11 embryos. Statistical significance was assessed relative to HSCs using a repeated-measures one-way analysis of variance (ANOVA) followed by either Dunnett’s multiple comparisons test (A, B) or Tukey’s multiple comparisons test (H, K), or using a two-tailed Student’s t-test to compare differences between fetal and adult cells (C-E and L-M); *P<0.05, **P<0.01, ***P<0.001.

Surprisingly, we found that adult erythroid progenitors synthesized ∼3.5-fold more protein than their fetal counterparts (Fig. 1E). We thus closely investigated protein synthesis in adult and fetal erythroid progenitors. Erythroid differentiation traverses five distinct stages (R1-R5) that can be identified based on CD71 and Ter119 expression^16^ (Fig. 1F,I). We examined protein synthesis within adult and fetal R1-R4 erythroid populations in vivo (Fig. 1G; R5 cells do not survive the OP-Puro procedure). In adult BM, protein synthesis increased ∼2-fold at the R1/R2 transition. It remained elevated through the R3 stage but declined ∼4-fold at the R4 stage (Fig. 1H). In fetal liver, protein synthesis again increased ∼2-fold at the R1/R2 transition, but significantly declined by the R3 stage and was ∼6-fold lower in R4 relative to R2 cells (Fig. 1J,K). Thus, fetal erythroid progenitors exhibit lower protein synthesis than their adult counterparts (Fig. 1L,M), and they attenuate protein synthesis earlier in the differentiation program.

Our findings raised the question of whether ontogeny-dependent changes in protein synthesis correlate with changes in sensitivity to ribosomal mutations. To test this, we utilized mice with a ribosomal protein L24 mutation (*Rpl24*^Bst/+^)^10^. Fetal and adult *Rpl24*^Bst/+^ HSCs exhibited 30-50% reductions in protein synthesis (Fig. 2A,B). Reduced protein synthesis did not significantly affect adult BM cellularity, HSC frequency or number (Fig. 2C,D; S1A). In contrast, *Rpl24*^Bst/+^ fetal liver cellularity was significantly reduced, and HSC frequency and number were increased compared to controls (Fig. 2E-G). *Rpl24*^Bst/+^ fetal HSCs proliferated at normal rates (Fig. 2H), consistent with the notion that protein synthesis and proliferation are partially decoupled in HSCs^3^. To test whether fetal HSC function is compromised by the *Rpl24*^Bst/+^ mutation, as we previously observed for adult HSCs (Fig. S1B,C)^1^, we transplanted 10 E15.5 HSCs from *Rpl24*^Bst/+^ or control mice (CD45.2^+^) with 3×10^5^ recipient-type BM cells into irradiated mice (CD45.1^+^) (Fig. S2A). The *Rpl24*^Bst^ mutation severely reduced HSC reconstituting activity (Fig. 2I,J; Fig. S2B-E), demonstrating that HSCs are exquisitely sensitive to reductions in protein synthesis, irrespective of whether they have high or low baseline protein synthesis.

**Figure 2.**
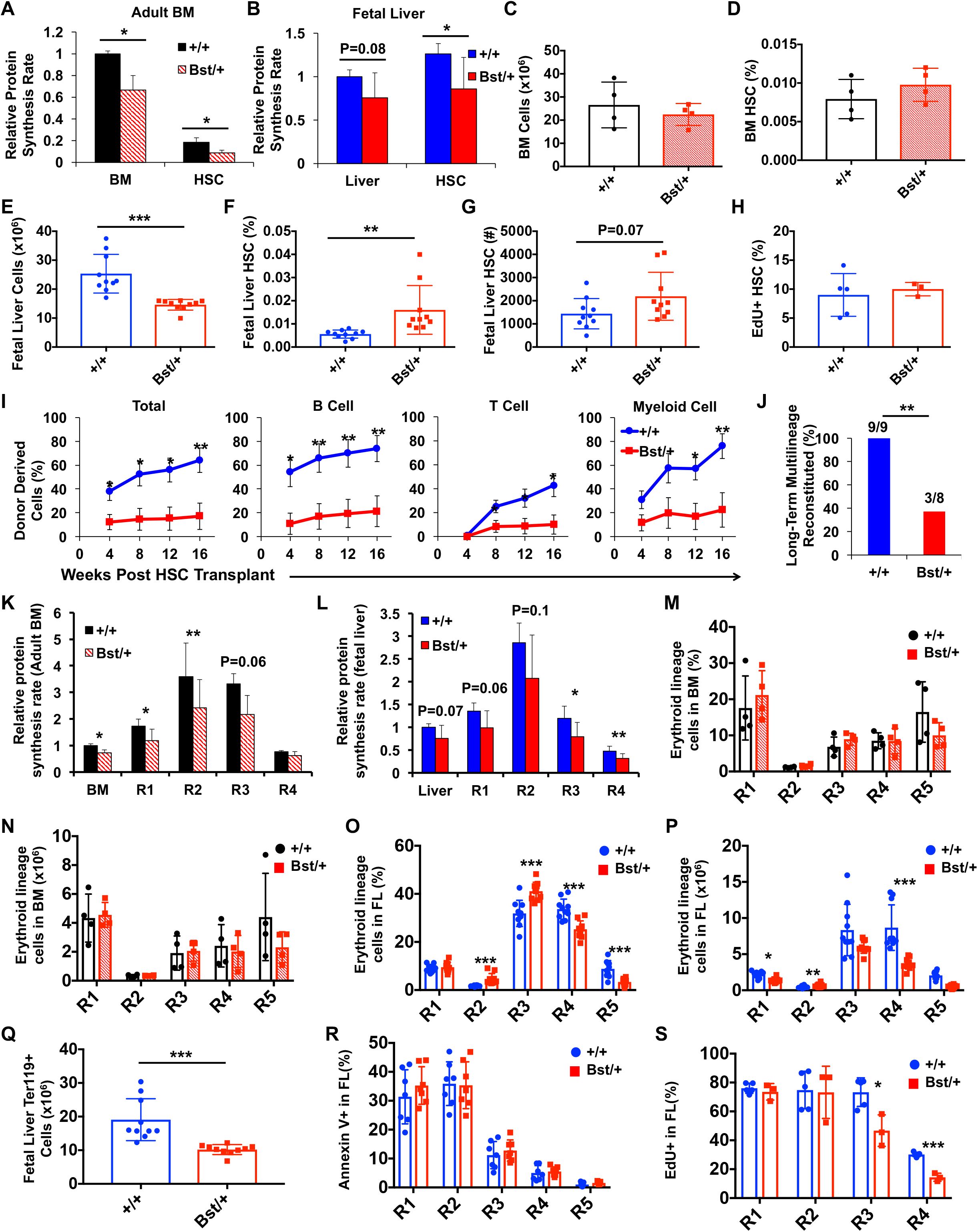
The *Rpl24*^Bst/+^ mutation impairs fetal and adult HSCs but only fetal erythroid lineage cells. (A,B) Relative protein synthesis rates in (A) unfractionated BM and HSCs in young adult *Rpl24*^Bst/+^ and control mice, and (B) unfractionated fetal liver cells and HSCs in E15.5 *Rpl24*^Bst/+^ and littermate control embryos. Values are normalized to control BM (N=4 mice/genotype) or control liver cells (N=12-13 embryos/genotype). (C) BM cellularity and (D) HSC frequency in young adult *Rpl24*^Bst/+^ and control mice (1 femur + 1 tibia/mouse; N=4 mice/genotype). (E) Fetal liver cellularity, (F) HSC frequency and (G) HSC number in E15.5 *Rpl24*^Bst/+^ and littermate control embryos (N=10 embryos/genotype). (H) Frequency of fetal liver HSCs that incorporated EdU after a 1-hour pulse in vivo (N=3-5 embryos/genotype). (I) Donor cell engraftment when 10 *Rpl24*^*Bst/+*^ (Bst/+) or littermate control (+/+) HSCs were transplanted with 3×10^5^ recipient-type young adult bone marrow cells into irradiated mice. Total hematopoietic, B-, T- and myeloid cell engraftment is shown 4, 8, 12 and 16 weeks after transplantation (N=8-9 recipients per genotype). Long-term reconstitution of individual recipients is shown in Fig. S2B-E. (J) Frequency of recipient mice in (I) that exhibited long-term (16-week) multilineage reconstitution (≥0.5% donor derived peripheral blood B-, T- and myeloid cells). (K,L) Relative protein synthesis in *Rpl24*^Bst/+^ (Bst/+) or control (+/+) erythroid progenitor cells in (K) young adult BM (N=4 mice/genotype) and (L) E15.5 fetal liver (N=12-13 embryos/genotype) in vivo. (M) Frequency and (N) absolute number of erythroid progenitors in *Rpl24*^Bst/+^ (Bst/+) or control (+/+) young adult BM (1 femur + 1 tibia/mouse; N=3-4 mice/genotype). (O) Frequency and (P) absolute number of erythroid progenitors in *Rpl24*^Bst/+^ (Bst/+) or control (+/+) E15.5 fetal liver (FL; N=10 embryos/genotype). (Q) Number of Ter119^+^ cells in *Rpl24*^Bst/+^ (Bst/+) or control (+/+) E15.5 fetal liver (N=10 embryos/genotype). (R) Frequency of erythroid progenitors that are Annexin V^+^ in *Rpl24*^Bst/+^ (Bst/+) or control (+/+) E15.5 fetal liver (N=7 embryos/genotype). (S) Frequency of *Rpl24*^Bst/+^ (Bst/+) or control (+/+) fetal liver erythroid progenitors that incorporated EdU after a 1-hour pulse in vivo (N=3-5 embryos/genotype). Data represent mean ± SD (A-H; K-S) or standard error of the mean (SEM; I). Statistical significance was assessed using a two-tailed Student’s t-test or a Fisher’s exact test (J); *P<0.05, **P<0.01, ***P<0.001.

We next tested whether erythroid progenitors are similarly sensitive to perturbations in protein synthesis across different stages of development. The *Rpl24*^Bst/+^ mutation reduced protein synthesis by 25-55% in fetal and adult erythroid progenitors (Fig. 2K,L). Adult *Rpl24*^Bst/+^ BM contained similar frequencies and numbers of erythroid progenitors as controls (Fig. 2M,N; S3A,B), and there was no change in erythroid progenitor apoptosis (Fig. S3C) or peripheral red blood cells^1^. In contrast, *Rpl24*^Bst/+^ fetal livers contained significantly more R2 and R3 erythroid progenitors and significantly fewer R4 and R5 cells compared to controls (Fig. 2O,P). Overall, *Rpl24*^Bst/+^ fetal livers contained ∼45% fewer Ter119^+^ erythroid cells than controls (Fig. 2Q). The depletion of fetal erythroid progenitors was not attributable to changes in apoptosis, as Annexin-V labeling was similar in *Rpl24*^Bst/+^ and control R1-R5 cells (Fig. 2R). However, *Rpl24*^Bst/+^ fetal R3 and R4 cells exhibited significantly reduced proliferation in vivo (Fig. 2S), which coincides with the stage at which fetal progenitors specifically exhibit a decline in protein synthesis. Low protein synthesis may thus hypersensitize differentiating fetal erythroid progenitors to ribosomal mutations by restricting proliferation and translation of essential mRNAs^17-19^.

This study reveals that protein synthesis is dynamically controlled in a cell-type- and developmental stage-specific manner in the hematopoietic system. Fetal HSCs maintain relatively high protein synthesis, yet they retain self-renewal capacity. Bile acids in the fetal liver protect HSCs by restricting activation of the unfolded protein response^20^, but a more complete understanding of how fetal HSCs cope with protein misfolding that accompanies high protein synthesis could yield interventions to improve proteostasis maintenance in adult HSCs^21^. Unlike HSCs, erythroid progenitors exhibit fetal-specific susceptibility to defects in ribosome biogenesis, suggesting that ribosomal mutations may disrupt erythropoiesis to a greater degree in early stages of life. Indeed, some Diamond Blackfan Anemia patients go into spontaneous remission as they age^22^. Further studies should resolve whether temporal changes in protein synthesis convey susceptibility to disease-relevant mutations^23^.

## Acknowledgements

Work in the Signer Laboratory is supported by the NIH/NIDDK (R01DK116951; R01DK124775), a Blood Cancer Discoveries Grant (8025-20) from the Leukemia & Lymphoma Society, The Mark Foundation for Cancer Research and the Paul G. Allen Frontiers Group, a Scholar Award from the American Society of Hematology, the Sanford Stem Cell Clinical Center, and the UCSD Moores Cancer Center which is supported by the NIH/NCI Specialized Cancer Center Support Grant (2P30CA023100). The LJI Flow Cytometry core facility is supported by the NIH Shared Instrumentation Grant Program (S10 RR027366). Work in the Magee Laboratory is supported by the NHLBI (R01HL136504 and R01HL152180), Alex’s Lemonade Stand Foundation (‘A’ Award), Hyundai Hope on Wheels, and the Children’s Discovery Institute of Washington University and St. Louis Children’s Hospital. We would like to thank Sean Morrison for support and resources and Kellie Machlus for advice.

## Authorship Contributions

J.A.M and R.A.J.S. conceived the project, performed HSC analyses and wrote the manuscript. R.A.J.S. performed protein synthesis and erythroid analyses.

