## Supplementary Information for "Developmental stage-specific changes in protein synthesis differentially sensitize hematopoietic stem cells and erythroid progenitors to impaired ribosome biogenesis"

### Supplementary Figure Legends

**Figure S1. Young adult *Rpl24*$^{Bst/+}$ HSCs have impaired long-term multilineage reconstituting activity.** (A) Absolute number of HSCs in young adult *Rpl24*$^{Bst/+}$ and control mice (1 femur + 1 tibia/mouse; N=4 mice/genotype). (B) Diagram of experimental strategy to test long-term multilineage reconstituting activity of young adult *Rpl24*$^{Bst/+}$ HSCs. (C) Donor cell engraftment when $5x10^5$ *Rpl24*$^{Bst/+}$ (Bst/+) or littermate control (+/+) BM cells were transplanted with $5x10^5$ recipient-type young adult BM cells into irradiated mice. Total hematopoietic, B-, T- and myeloid cell engraftment is shown 4, 8, 12 and 16 weeks after transplantation (N=5 recipients/genotype). Data represent mean ± SD (A) or SEM (C). Statistical significance was assessed using a two-tailed Student's t-test (*P<0.05, **P<0.01, ***P<0.001).

**Figure S2. Fetal liver *Rpl24*$^{Bst/+}$ HSCs have impaired long-term multilineage reconstituting activity.** (A) Diagram of experimental strategy to test long-term multilineage reconstituting activity of fetal liver *Rpl24*$^{Bst/+}$ HSCs. (B-E) Long-term (16-week) donor (B) hematopoietic, (C) B-, (D) T- and (E) myeloid cell engraftment in the peripheral blood of individual recipient mice from transplants in Fig. 2I. Data represent mean ± SEM. Statistical significance was assessed using a two-tailed Student's t-test (*P<0.05, **P<0.01).

**Figure S3. The *Rpl24*$^{Bst/+}$ mutation impairs fetal but not adult erythroid differentiation.** (A) Representative flow cytometry plots of R1-R5 erythroid lineage cells in *Rpl24*$^{Bst/+}$ (Bst/+) or control (+/+) young adult BM. (B) Number of Ter119$^+$ cells in *Rpl24*$^{Bst/+}$ (Bst/+) or control (+/+) young adult *Rpl24*$^{Bst/+}$ (Bst/+) or control (+/+) BM (1 femur + 1 tibia/mouse; n=4 mice/genotype). (C) Frequency of erythroid progenitors that are Annexin V$^+$ in young adult *Rpl24*$^{Bst/+}$ (Bst/+) or control (+/+) BM (N=3 mice/genotype). (D) Representative flow cytometry plots of R1-R5 erythroid lineage cells in

*Rpl24*$^{Bst/+}$ (Bst/+) or control (+/+) E15.5 fetal liver. Data represent mean ± SD. Statistical significance was assessed using a two-tailed Student's t-test.

### SUPPLEMENTARY FIGURE S1

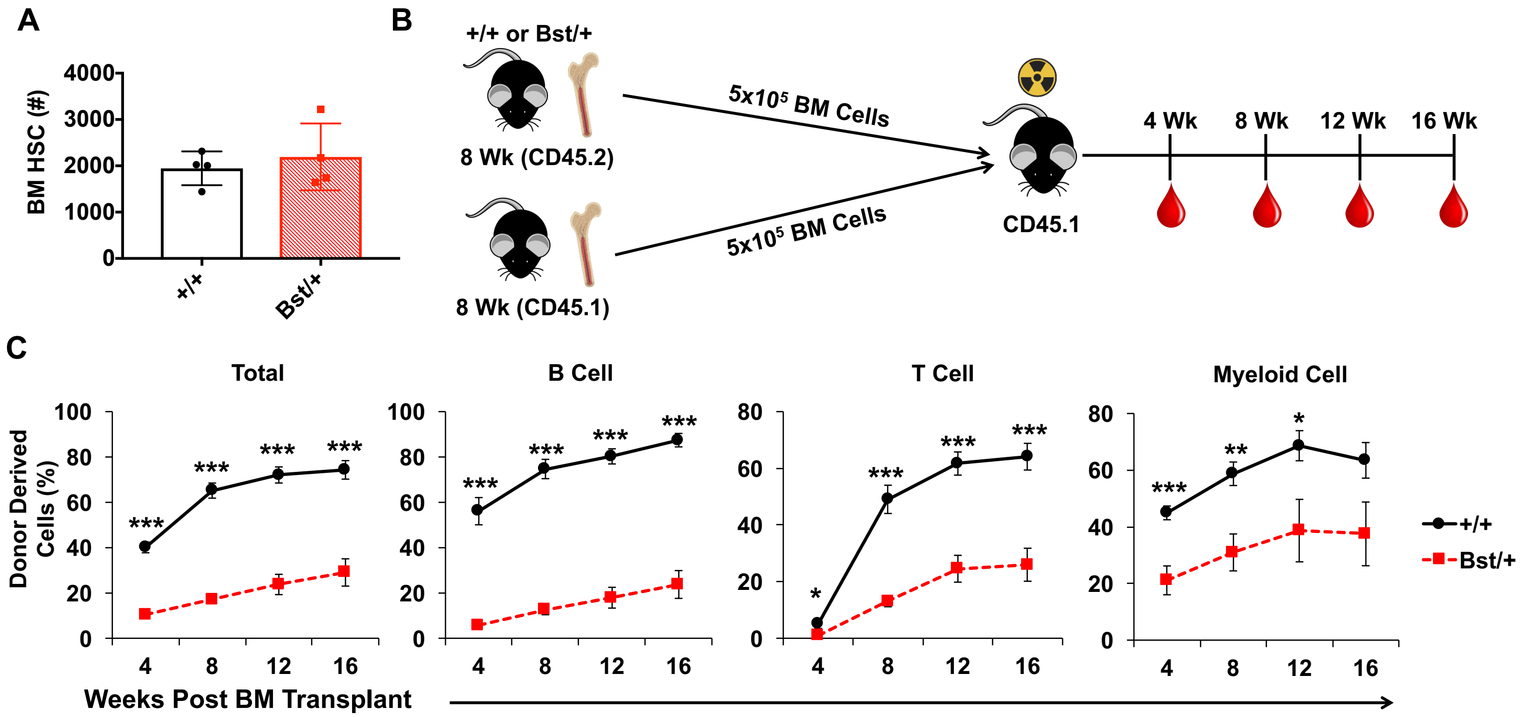

### SUPPLEMENTARY FIGURE S2

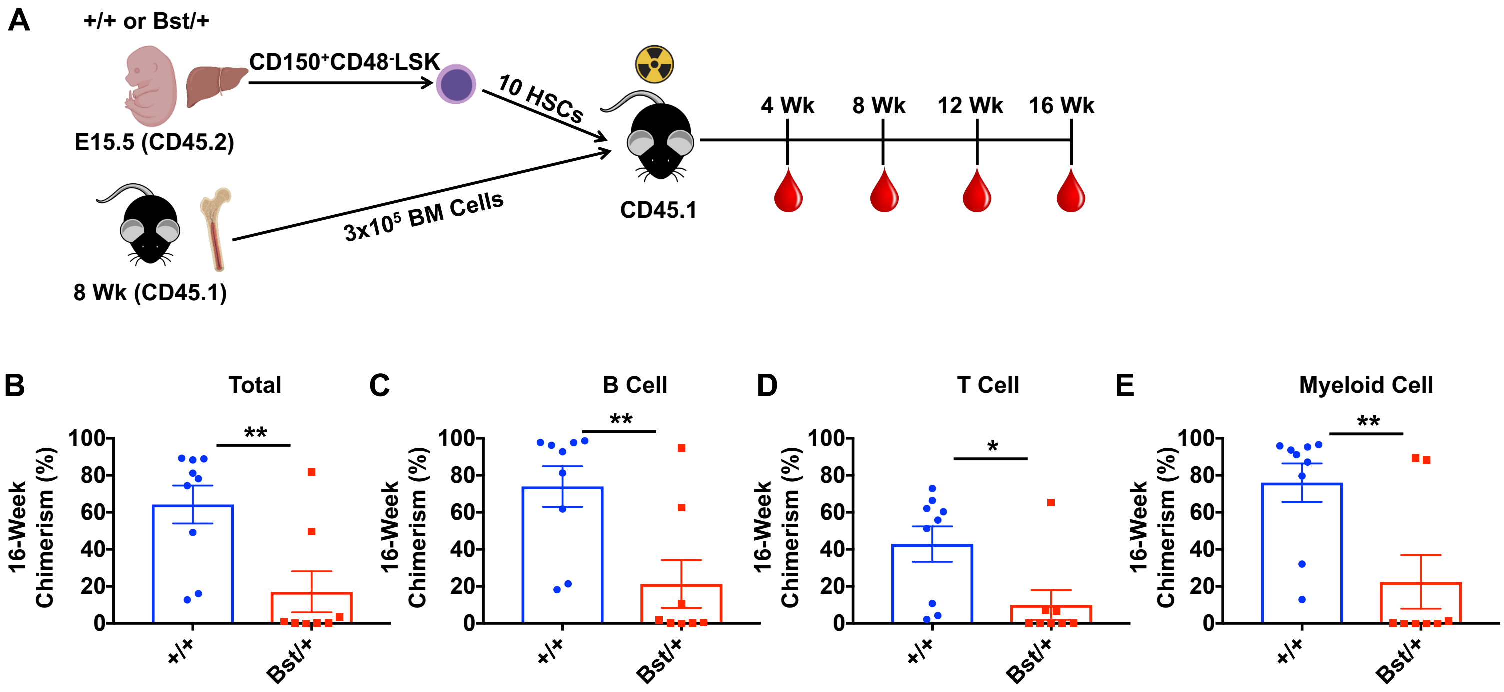

### SUPPLEMENTARY FIGURE S3

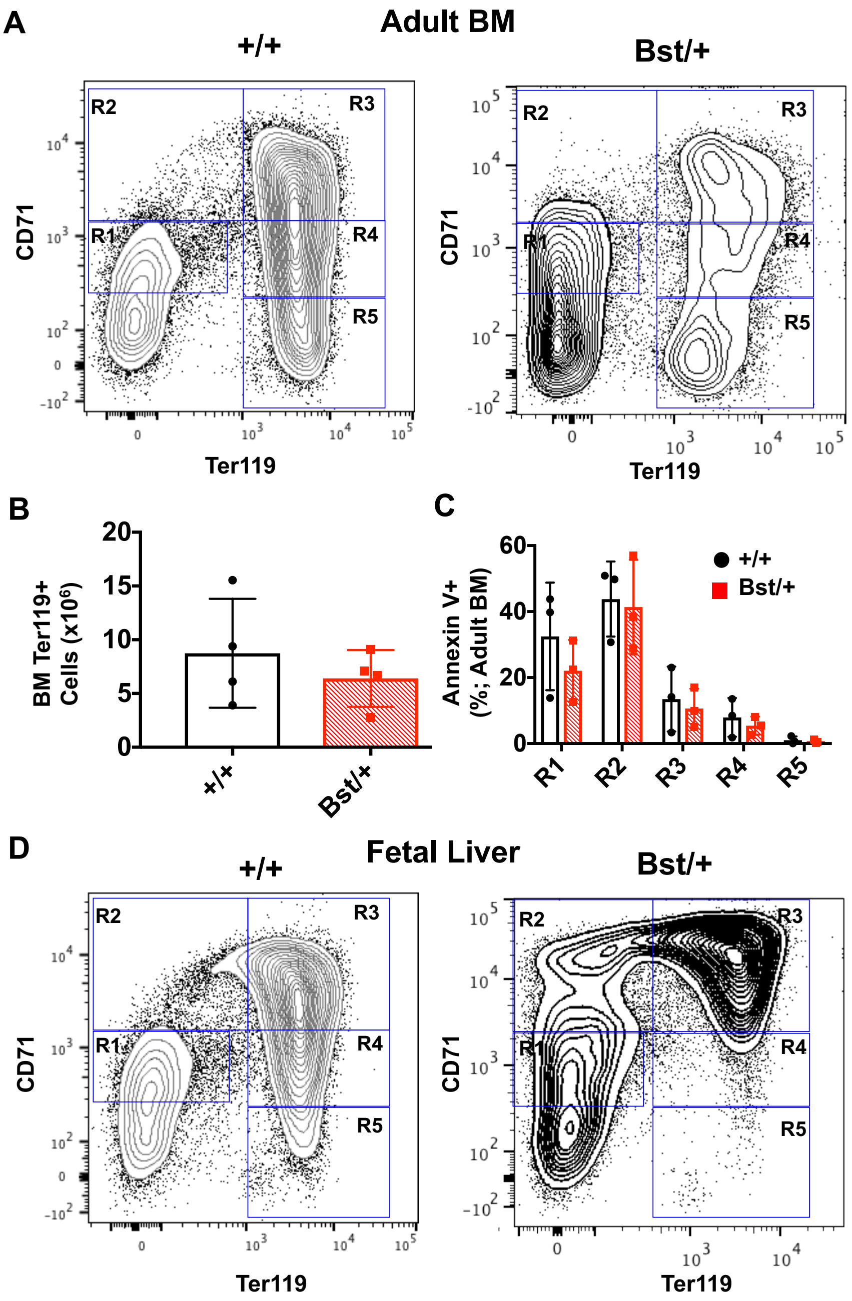
